## Supplemental Table and Figures for "Hemozoin produced by mammals confers heme tolerance"

### Supplemental Table 1

| Mutant Name | Strain | Mutation | Sequence |
|---|---|---|---|
| **M1** | C57BL/6Jx129/SvJ | 7bp deletion | …GACGGTGGTCTACCG•••••••GGGACTGCGGCGAT… |
| **M2** | C57BL/6Jx129/SvJ | 7bp deletion | …GACGGTGGTCTACCGA•••••••GGACTGCGGCGAT… |
| **M3** | C57BL/6Jx129/SvJ | 18bp deletion | …GACGG••••••••••••••••••GGACTGCGGCGAT… |
| **M4** | C57BL/6Jx129/SvJ | 1bp insertion | …GACGGTGGTCTACCGACAAACCGGGGACTGCGGCGAT… |
| **M5** | C57BL/6Jx129/SvJ | 1bp deletion | …GACGGTGGTCTACCGACA•CCGGGGACTGCGGCGAT… |
| **M6** | C57BL/6Jx129/SvJ | 2bp deletion | …GACGGTGGTCTACCGA••ACCGGGGACTGCGGCGAT… |
| **M7** | C57BL/6Jx129/SvJ | 11bp deletion | …GACGGTGG•••••••••••CCGGGGACTGCGGCGAT… |
| **Wildtype** | | | …GACGGTGGTCTACCGACAACCGGGGACTGCGGCGAT… |

#### Supplemental Table 2

| | Standard diet | | 2ppm diet | | n |
|---|---|---|---|---|---|
| | +/+ | -/- | +/+ | -/- | |
| **Serum iron (μg/dl)** | 113.5±11.72[a] | 129.3±11.34[a] | 66.32±13.62[b] | 65.63±17.09[b] | 8-15 |
| **TIBC (μg/dl)** | 348.4±32.1[a] | 341.8±11.62[a] | 451.8±17.08[b] | 459±19.92[b] | 8-15 |
| **Tf saturation (%)** | 36.05±4.452[a] | 37.35±2.601[a] | 14.3±2.975[b] | 14.6±4.034[b] | 8-15 |
| **Serum ferritin (ng/ml)** | 369.1±57.02[a] | 585.9±58.62[b] | 284.7±63.8[a] | 492±63.77[b] | 9-15 |

Values with different letters are statistically different from each other[a,b]

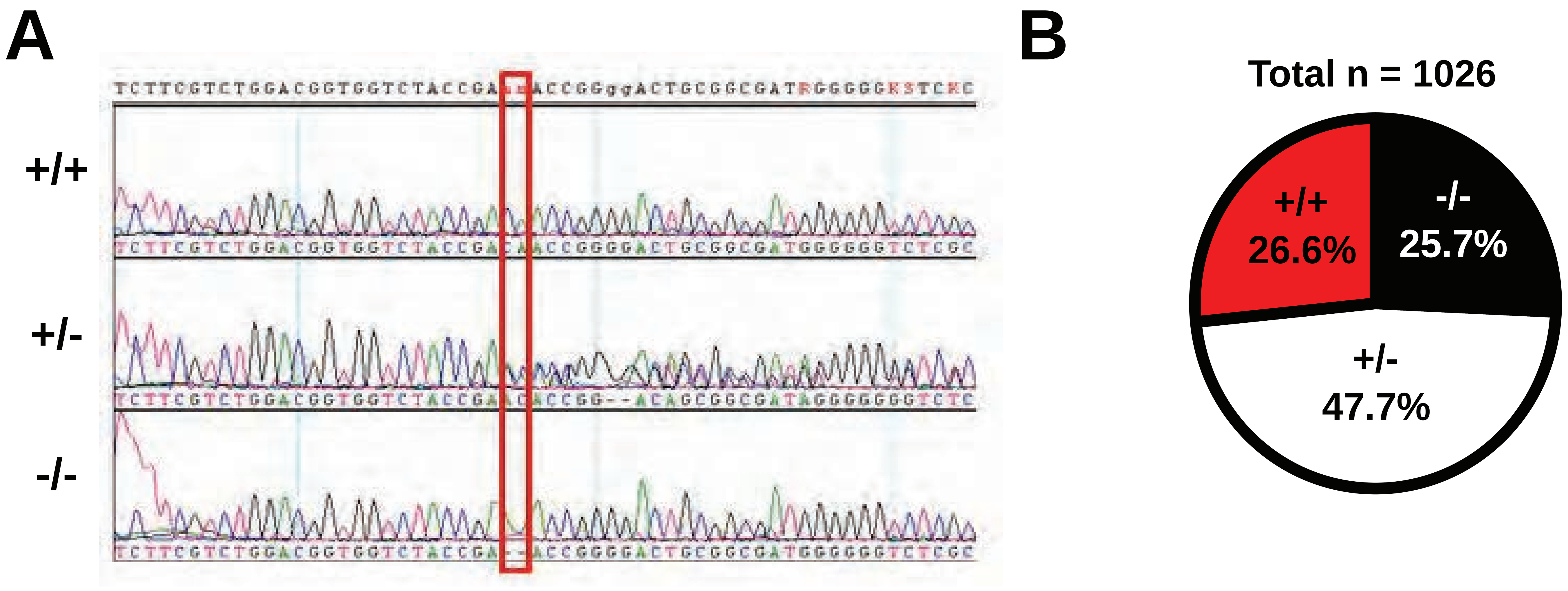


**Supplemental figure 1**

**A)** Chromatogram of genomic DNA sequencing from mice indicating 2 basepair deletion. **B)** Mendelian distribution of P21 pups derived from *HRG1*$^{+/-}$ intercrosses.

### Supplemental Table 3

| Spleen | | | | | | | |
|---|---|---|---|---|---|---|---|
| Standard | | | | 2ppm | | | |
| Upregulated | p-value | Downregulated | p-value | Upregulated | p-value | Downregulated | p-value |
| Pcbp4 | 0.020129 | Slc48a1 | 0.023006 | Heph | 0.00032 | | |
| | | Cp | 0.040584 | Bmp6 | 0.00041 | | |
| | | | | Hfe2 | 0.000626 | | |
| | | | | Neo1 | 0.001396 | | |
| | | | | Il6 | 0.003406 | | |
| | | | | Pcbp4 | 0.003501 | | |
| | | | | Il10 | 0.004212 | | |
| | | | | Stat6 | 0.006443 | | |
| | | | | Hpx | 0.007156 | | |
| | | | | Steap1 | 0.00791 | | |
| | | | | Tmprss6 | 0.007936 | | |
| | | | | Ifna4 | 0.008373 | | |
| | | | | Saa1 | 0.008779 | | |
| | | | | Nox4 | 0.008779 | | |
| | | | | Epo | 0.008779 | | |
| | | | | Gast | 0.008779 | | |
| | | | | Il22 | 0.008779 | | |
| | | | | Hamp | 0.008779 | | |
| | | | | Nox3 | 0.008779 | | |
| | | | | Steap2 | 0.00879 | | |
| | | | | Nox1 | 0.008795 | | |
| | | | | Ifnb1 | 0.010902 | | |
| | | | | Asb11 | 0.011532 | | |
| | | | | Fbxl5 | 0.020012 | | |
| | | | | Tlr3 | 0.027149 | | |
| | | | | Pcbp3 | 0.027975 | | |
| | | | | Hfe | 0.037129 | | |
| | | | | Ifna2 | 0.041967 | | |
| | | | | Hp | 0.047671 | | |
| | | | | Cp | 0.050688 | | |
| | | | | Slc25a38 | 0.051437 | | |
| | | | | Slc38a1 | 0.054551 | | |
| | | | | Tlr7 | 0.05663 | | |
| | | | | Mfsd7b | 0.056827 | | |
| | | | | Hmox1 | 0.057685 | | |

| Liver | | | | | | | |
|---|---|---|---|---|---|---|---|
| Standard | | | | 2ppm | | | |
| Upregulated | p-value | Downregulated | p-value | Upregulated | p-value | Downregulated | p-value |
| Ifna2 | 0.010002 | Slc48a1 | 0.004204 | | | Iscu | 0.004343 |
| Cybrd1 | 0.010447 | | | | | Slc48a1 | 0.004492 |
| | | | | | | Ncoa4 | 0.010102 |
| | | | | | | Slc11a2 | 0.01464 |
| | | | | | | Pcbp2 | 0.040291 |

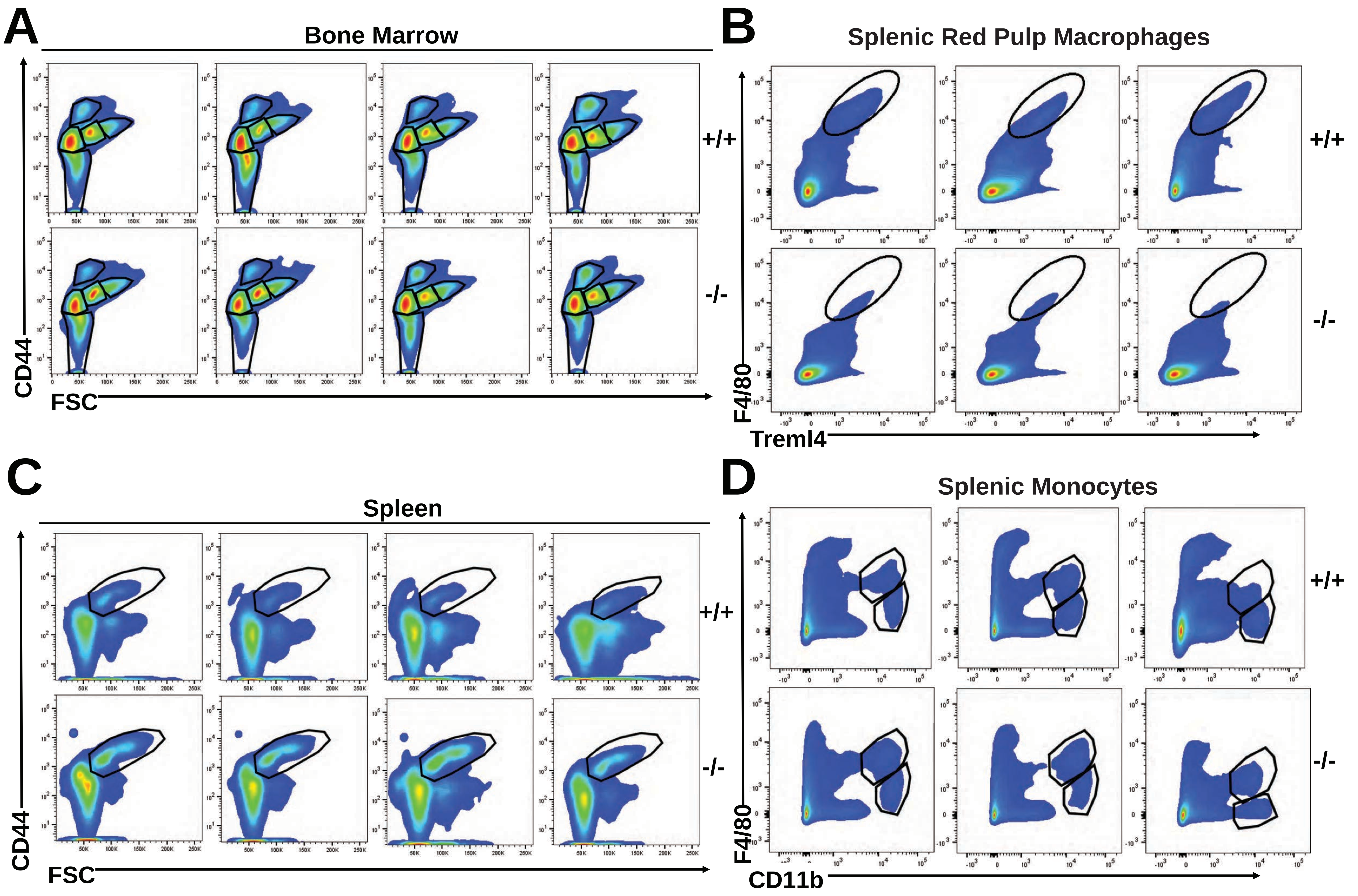


**Supplemental figure 2**

**A)** Gating of bone marrow Ter-119⁺ subpopulations. **B)** Gating of splenic Ter-119⁺ population II+III. **C-D)** Gating of splenic RPMs and monocytes. Individual plots shown are representative of all mice analyzed per group.

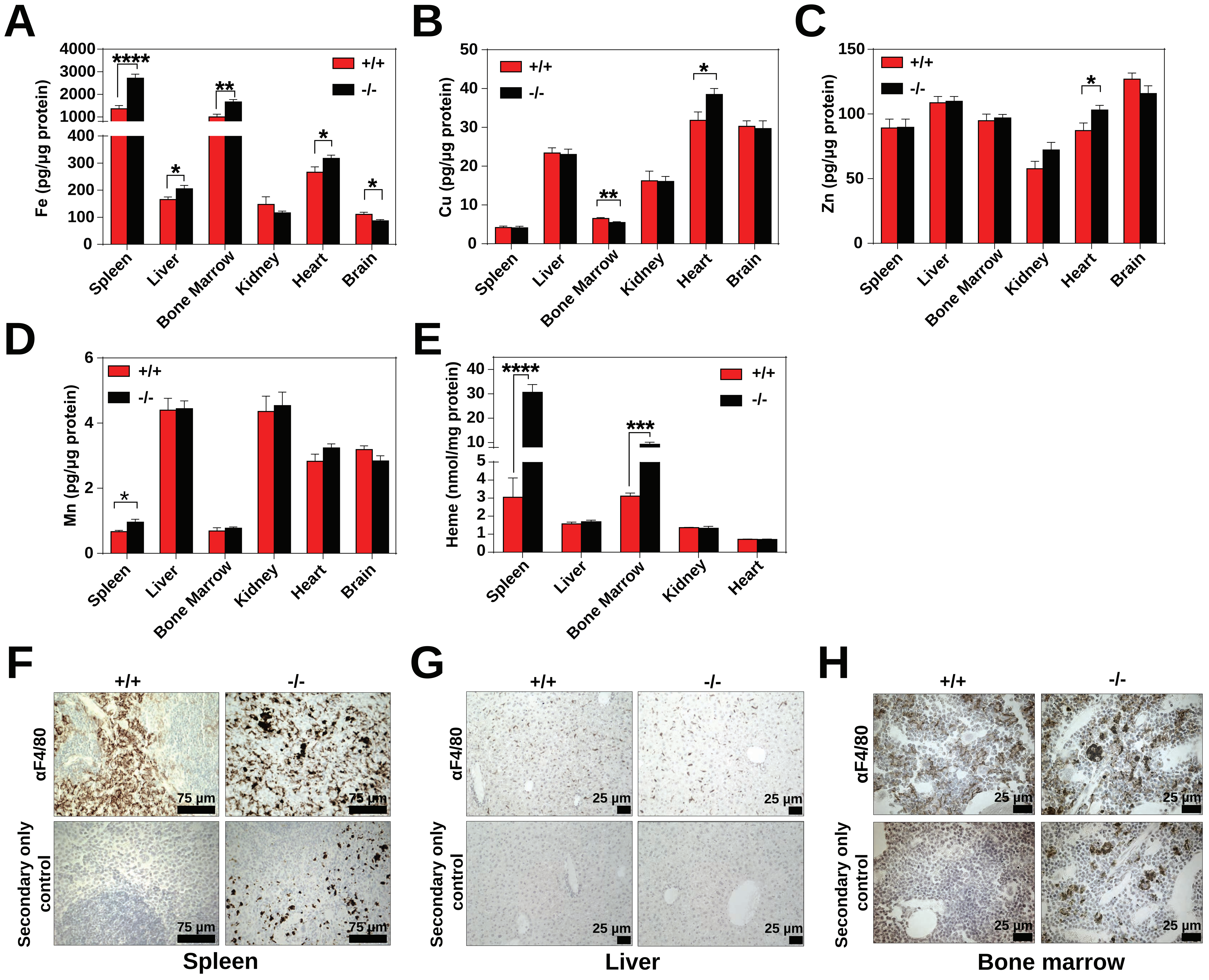


**Supplemental Figure 3**
**A-E)** Quantification of tissue iron (Fe), copper (Cu), zinc (Zn), manganese (Mn) and heme (n=6-17). Heme was undetectable in brain tissue. F4/80 immunohistochemistry of spleen **(F)**, liver **(G)** and bone marrow **(H)** sections. *$p$<0.05; **$p$<0.01; ***$p$<0.001; ****$p$<0.0001.

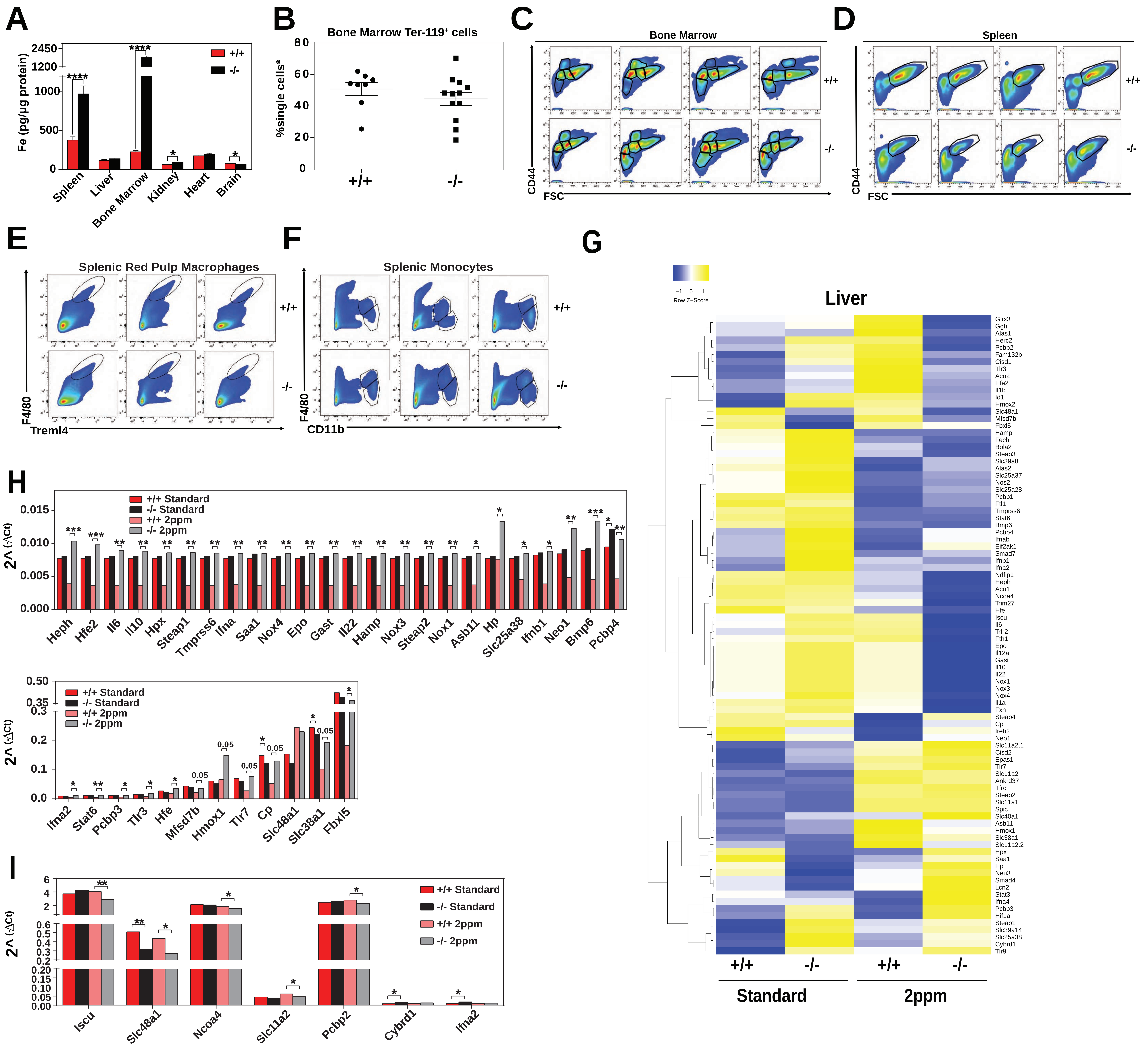


**Supplemental Figure 4**

**A)** Quantification of tissue iron (Fe) (n=6-17). **B)** Quantification of total Ter-119+ cells represented as a percentage of all single cells analyzed in the bone marrow. The %single cells* on the y-axis denote single cells that are negative for CD4/8/41, B220 and Gr-1. At least 100,000 single cells were analyzed per sample. Gating of **C)** splenic Ter-119+ populations II+III and **D)** bone marrow Ter-119+ subpopulations. **E, F)** Gating of splenic RPMs and monocytes. Individual plots shown are representative of all mice analyzed per group. **G)** Gene expression heat map of 90 iron metabolism genes in livers from mice on standard or low iron (2 ppm) diet. Pearson correlation was used for comparison; average linkage (n=9 per group, per genotype). Gene expression by qPCR of iron metabolism genes in spleens **(H,** top and bottom panel separated by expression level) and livers **(I)** of *HRG1*$^{+/+}$ and *HRG1*$^{-/-}$ mice on indicated diets (n=9 mice per group). Gene expression was calculated as described in the experimental procedures section. *$p$<0.05; **$p$<0.01; ***$p$<0.001; ****$p$<0.0001.

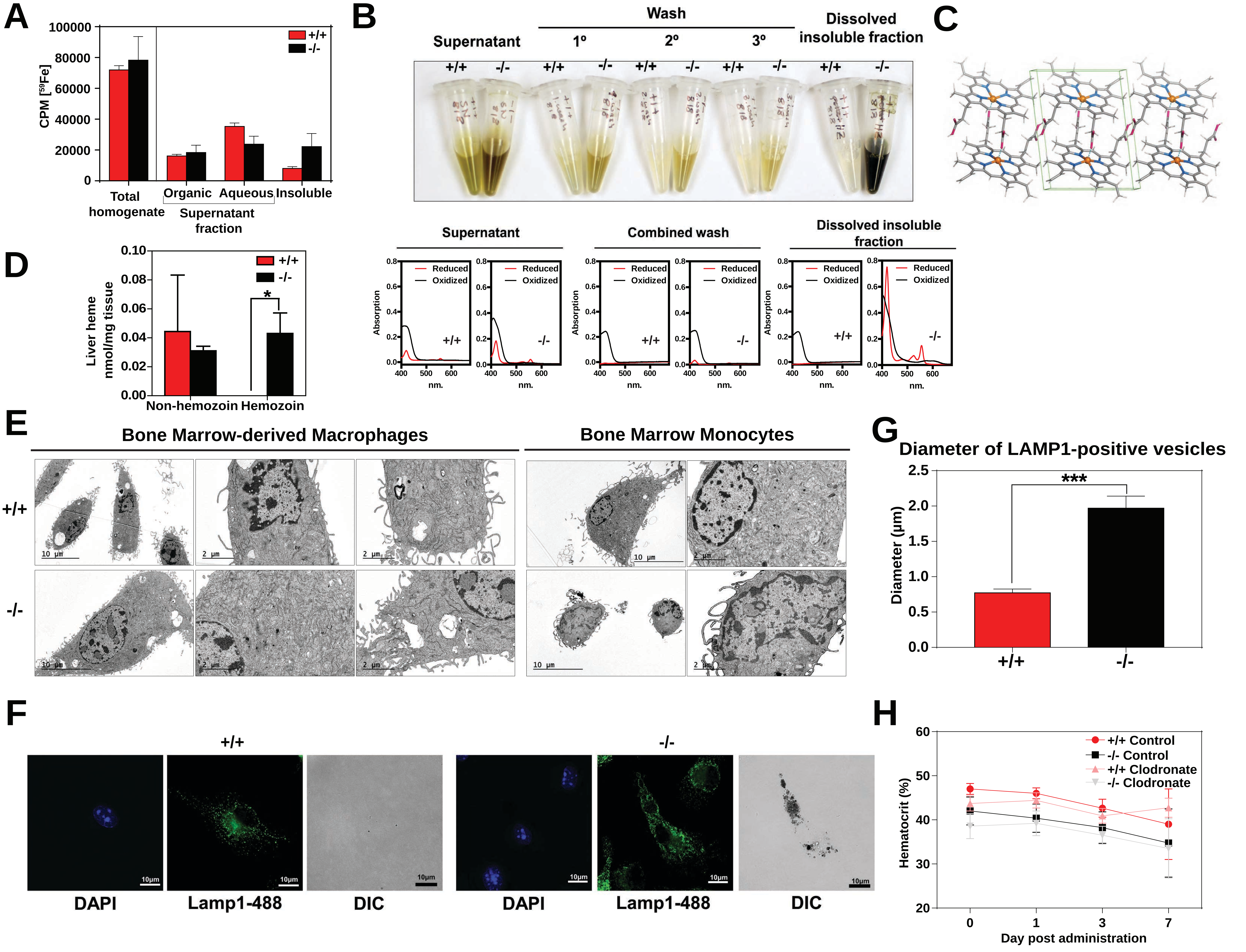


**Supplemental Figure 5**

**A)** $^{59}$Fe retained in differentially extracted fractions of the liver at 96 h, represented as counts per min. Total homogenate: homogenized and proteinase-treated whole spleen; Organic: ethyl acetate extractable [$^{59}$Fe]heme; Aqueous: ethyl acetate non-extractable $^{59}$Fe; Insoluble fraction: proteinase-insoluble fraction containing $^{59}$Fe. n=4-6. **B)** Images of different fractions of spleen homogenates, accompanied by spectrophotometric measurements of each fraction (lower panel). **C)** Chemical structure (rotated view) of hemozoin from *HRG1*$^{-/-}$ mice. **D)** Quantification of liver heme by spectrophotometric measurements (n=3). **E)** TEM of in vitro-differentiated bone marrow macrophages and isolated bone marrow monocytes. At least 3 cells were imaged per genotype. **F)** LAMP1 immunofluorescence on F4/80$^{+}$ bone marrow macrophages from *HRG1*$^{+/+}$ and *HRG1*$^{-/-}$ mice. **G)** Diameter of LAMP1-positive vesicles observed in F4/80$^{+}$ bone marrow macrophages. In both *HRG1*$^{+/+}$ and *HRG1*$^{-/-}$ cells, the diameters of LAMP1-positive structures with distinguishable boundaries were quantified manually by ImageJ software analysis. *HRG1*$^{+/+}$*:* 4 cells, 3-5 structures per cell; *HRG1*$^{-/-}$*:* 11 cells, 3-12 structures per cell. **(H)** Hematocrits of iron-deficient *HRG1*$^{+/+}$ and *HRG1*$^{-/-}$ mice treated with control or clodronate liposomes (n=6-14). *$p$<0.05, ***$p$<0.001.

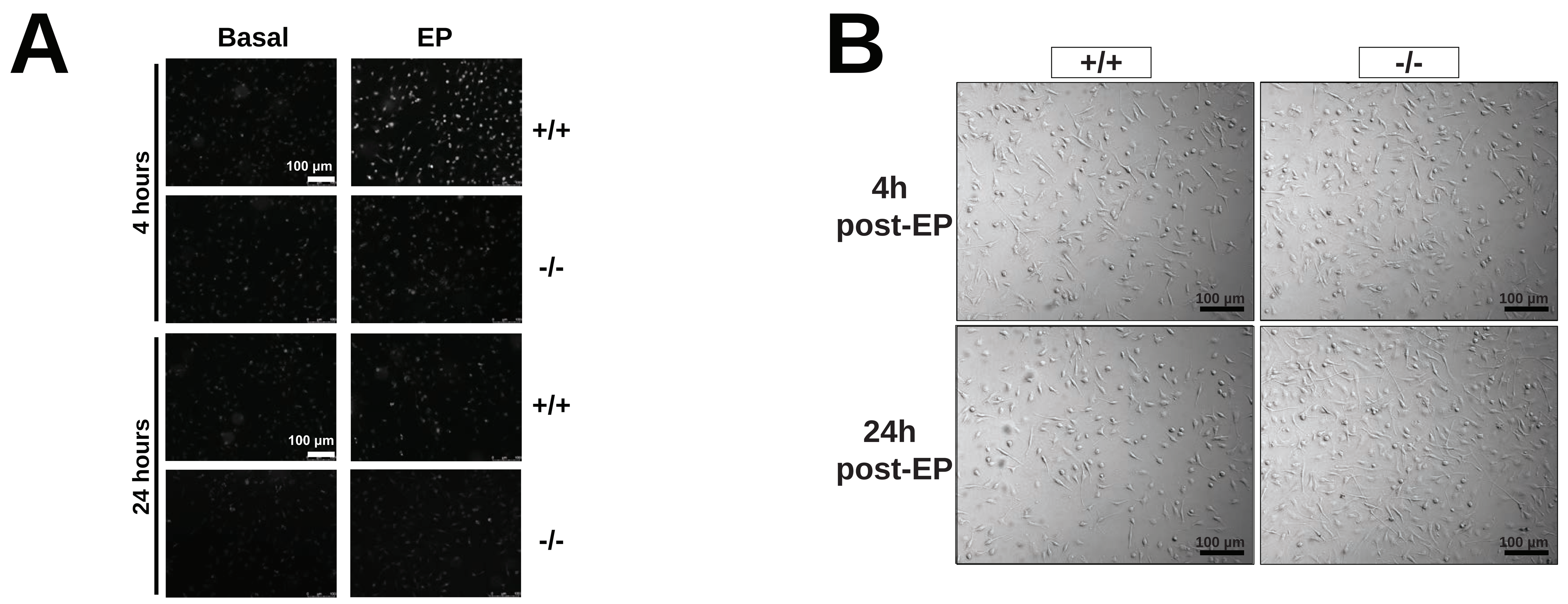


C

*HMOX1*+/-; *HRG1*+/+ x *HMOX1*+/-; *HRG1*+/+

| Genotype | Observed number | Expected number | Observed percentage |
|---|---|---|---|
| HMOX1+/+;HRG1+/+ | 64 | 64 | 100% |
| HMOX1+/-;HRG1+/+ | 133 | 128 | 104% |
| HMOX1-/-;HRG1+/+ | 9 | 64 | 14% |

*HMOX1*+/-; *HRG1*-/- x *HMOX1*+/-; *HRG1*-/-

| Genotype | Observed number | Expected number | Observed percentage |
|---|---|---|---|
| HMOX1+/+;HRG1-/- | 86 | 86 | 100% |
| HMOX1+/-;HRG1-/- | 108 | 172 | 63% |
| HMOX1-/-;HRG1-/- | 9 | 86 | 10% |

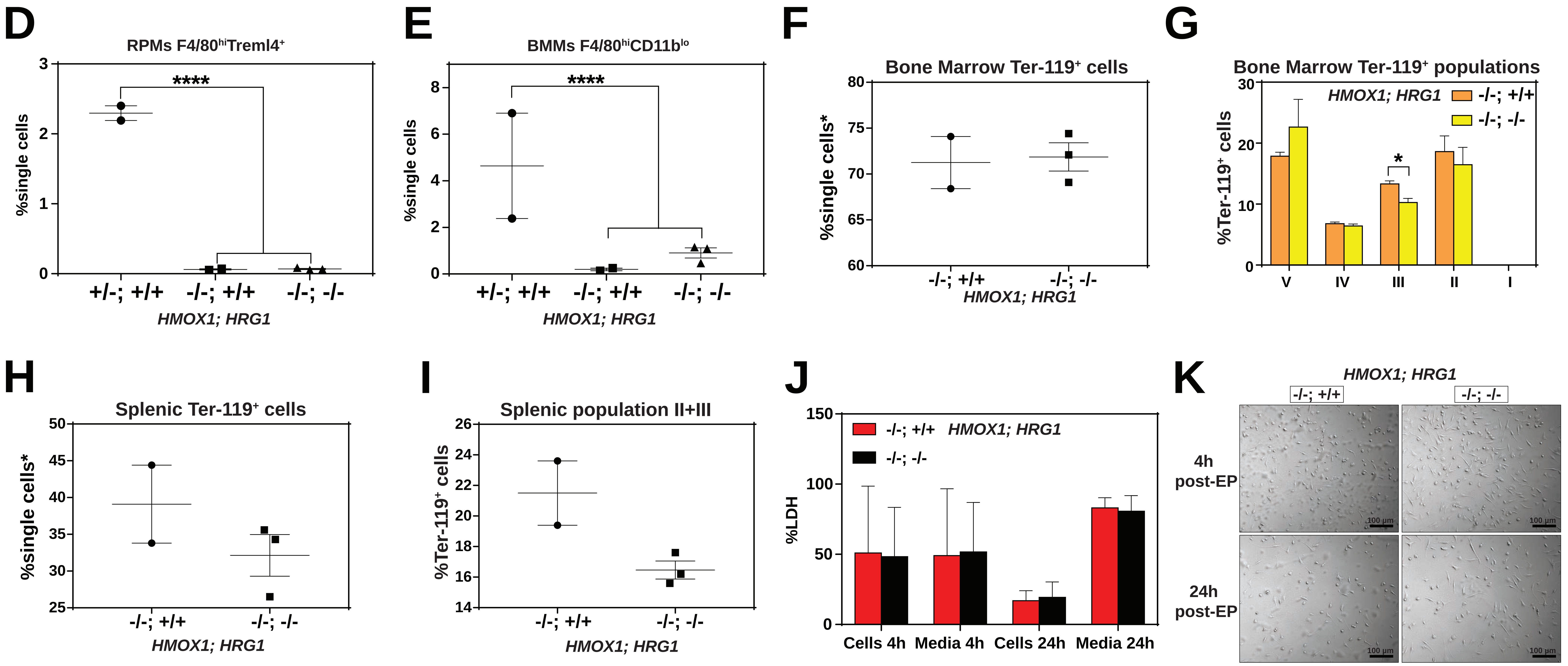


**Supplemental Figure 6**

**(A)** Representative images of *HRG1*$^{+/+}$ and *HRG1*$^{-/-}$ BMDMs post-EP. **B)** Representative images of intracellular reactive oxygen species (ROS) in *HRG1*$^{+/+}$ and *HRG1*$^{-/-}$ BMDMs at the indicated timepoints post-treatments. **C)** Observed and expected numbers of pups born of the indicated genotypes for *HMOX1*$^{+/-}$ intercrosses, either on a *HRG1*$^{+/+}$ or *HRG1*$^{-/-}$ background. Quantification of splenic RPMs **(D)** and bone marrow macrophages (BMMs) **(E)** in *HMOX1*$^{-/-}$*HRG1*$^{+/+}$ and *HMOX1*$^{-/-}$*HRG1*$^{-/-}$ mice by flow cytometry. **F)** Quantification of total Ter-119$^{+}$ cells in the bone marrows of *HMOX1*$^{-/-}$*HRG1*$^{+/+}$ and *HMOX1*$^{-/-}$*HRG1*$^{-/-}$ mice. **G)** Quantification of subpopulations of Ter-119$^{+}$ cells represented as a percentage of total Ter-119$^{+}$ cells in the bone marrow. **H)** Quantification of total Ter-119$^{+}$ cells in the spleen. The % single cells* on the y-axis denote single cells that are negative for CD4/8/41, B220 and Gr-1. **I)** Quantification of populations II and III of Ter-119$^{+}$ cells represented as a percentage of total Ter-119$^{+}$ cells in the spleen. At least 100,000 single cells were analyzed per sample. Each dot represents one mouse. **J)** LDH content in cells and media of *HMOX1*$^{-/-}$*HRG1*$^{+/+}$ and *HMOX1*$^{-/-}$*HRG1*$^{-/-}$ BMDMs post-erythrophagocytosis (EP), represented as percentage of total LDH in cells and media. **K)** Representative images of *HMOX1*$^{-/-}$*HRG1*$^{+/+}$ and *HMOX1*$^{-/-}$*HRG1*$^{-/-}$ BMDMs post-EP.
